# The Landscape of Sex- and APOE Genotype-Specific Transcriptional Changes in Alzheimer’s Disease at the Single Cell Level

**DOI:** 10.1101/2024.12.01.626234

**Authors:** Gefei Yu, Abigail Thorpe, Qi Zeng, Erming Wang, Dongming Cai, Minghui Wang, Bin Zhang

## Abstract

Alzheimer’s disease (AD) is the most common form of dementia, with approximately two-thirds of AD patients are females. Basic and clinical research studies show evidence supporting sex-specific differences contributing to the complexity of AD. There is also strong evidence supporting sex-specific interaction between the primary genetic risk factor of AD, *APOE4* and AD-associated neurodegenerative processes. Recent studies by us and others have identified sex and/or APOE4 specific differentially expressed genes in AD based on the bulk tissue RNA-sequencing data of postmortem human brain samples in AD. However, there lacks a comprehensive investigation of the interplay between sex and APOE genotypes at the single cell level. In the current study, we systematically explore sex and APOE genotype differences in single cell transcriptomics in AD. Our work provides a comprehensive overview of sex and APOE genotype-specific transcriptomic changes across 54 high-resolution cell types in AD and highlights individual genes and brain cell types that show significant differences between sexes and APOE genotypes. This study lays the groundwork for exploring the complex molecular mechanisms of AD and will inform the development of effective sex- and APOE-stratified interventions for AD.

## INTRODUCTION

Alzheimer’s disease (AD) is the most prevalent neurodegenerative disorder of aging ^1^, with approximately two-thirds of AD patients are females ^2^. There are conflicting literature reports regarding AD incidents among males and females with some suggesting a higher AD incidence in females than males and others suggesting no differences ^3-7^. Evidence from basic and clinical research studies supports sex-specific differences contributing to the complexity of AD. For example, sex-biased changes in brain structures and connectome have been demonstrated in AD subjects using various neuroimaging modalities. It was found that the rates of brain atrophy in females were 1-1.5% faster than those in males, most prominently seen in the MCI group with a 1% increase in atrophic rates and 2% in ventricular expansion ^8^. Sex-specific brain region atrophy have also been reported ^9^. Moreover, a significantly higher reduction of hippocampal integrity in female AD subjects was reported when compared to male subjects with Magnetic Resonance Imaging studies of the Minimal Interval Resonance Imaging in AD database ^10^.

There is strong evidence supporting sex-specific interaction between the primary genetic risk factor of AD, *APOE4* ^11^ and AD-associated neurodegenerative processes ^12, 13^. The sex-specific association between *APOE4* and tau has been reported ^14, 15^. Furthermore, studies suggest a stronger association of *APOE4* with cortical thinning, volume loss, brain connectivity and hypo-metabolism in AD females than males ^16-18^. Moreover, stronger associations of *APOE4* with cognitive impairment, memory decline, and neuropsychiatric symptoms of AD in females were reported ^19,20^. On the other hand, male-specific associations with cerebral amyloid angiopathy (CAA) and microbleed in AD have been reported ^21^. Even so, little has been done to understand the interplay between *APOE* and sex in ageing and AD.

Recently, we systematically identified sex and APOE4 specific differentially expressed genes in AD^22^ based on the bulk tissue RNA-sequencing data of postmortem human brain samples from the Mount Sinai Brain Bank (MSBB) cohort ^23^ and the Religious Orders Study and Rush Memory and Aging Project (ROSMAP) cohort ^24^. We further developed sex-specific gene network models of AD to predict sex specific driver genes of AD and subsequently validated *LRP10* (lipoprotein receptor related protein 10) as a female-specific driver of AD pathogenesis^22^.

In the current study, we further investigate the interplay between sex and APOE in AD at single cell level through differential gene expression analysis of single-nucleus RNA sequencing data. The findings not only provide a comprehensive overview of sex and APOE genotype-specific transcriptomic changes across 54 high-resolution cell types in AD but also highlight individual genes and brain cell types that show significant differences between sexes and APOE genotypes. This study lays the groundwork for exploring the complex molecular mechanisms of AD and will inform the development of effective sex- and APOE-stratified interventions for AD.

## RESULTS

We leveraged a single-nucleus RNA sequencing (snRNA-seq) dataset which was generated from postmortem brain tissues of subjects with or without dementia by Mathys et al.^25^. The dataset profiled 2,359,994 nuclei from 427 prefrontal cortex brain samples, including 215 female and 212 male donors. Disease status was categorized using the final consensus cognitive diagnosis score (cogdx), where 1 = no cognitive impairment (NCI), 2 = mild cognitive impairment (MCI), 4 = Alzheimer’s disease (AD), and 3, 5, 6 were categorized as other dementia. For this study, we focused solely on NCI (n = 146) and AD (n = 144) samples. To investigate the impact of *APOE* genotypes, we grouped participants as follows: *APOE* e22 and e23 were categorized as *APOE* e2 carriers (e2x; n = 58), e33 remained as e33 (n = 252), and e34 and e44 were classified as *APOE* e4 carriers (e4x; n=105). *APOE* e24 carriers were excluded from the analysis due to the controversial nature of their association with disease risk.

We utilized the 54 cell clusters identified in the original study^25^: 14 excitatory neuron clusters, 25 inhibitory neuron clusters, 1 oligodendrocyte cluster, 1 oligodendrocyte precursor cell cluster, 3 astrocyte cell clusters, 3 microglia cell clusters, 1 central nervous system (CNS)-associated macrophage (CAM) cluster, 1 T cell cluster, 1 endothelial cells (End) cluster, 1 smooth muscle cell (SMC) cluster, 2 fibroblasts (Fib) clusters, and 1 pericytes (Per) cell cluster.

### Cell type specific gene expression changes in AD in each sex-APOE subgroup

To investigate the transcriptional changes associated with AD in each *APOE* genotype (e2x, e33, and e4x) or sex group (female (F) and male (M)), we performed cell-cluster-specific differential gene expression (DEG) analysis across six stratified comparisons. For each sex-*APOE* group (F_e2x, F_e33, F_e4x, M_e2x, M_e33, and M_e4x), we compared gene expression profiles of the individuals with AD with those with no cognitive impairment (NCI). This stratification allowed us to capture genotype- and sex-specific transcriptional alterations in AD in each cell cluster, offering a detailed view of how these factors shape the molecular landscape in AD. The numbers of up-regulated and down-regulated DEGs identified in these comparisons are shown in **Figure 1**. The distribution of DEGs across cell clusters revealed distinct patterns. Excitatory neurons exhibited a substantial number of DEGs in almost all sex-*APOE* groups. Notably, the AD males with *APOE* e2 carriers (M_e2x) had the highest number of DEGs compared to other sex-*APOE* groups, particularly in the excitatory neuron (Exc), inhibitory neuron (Inh), OPC, and microglia (Mic) clusters. This suggests a potentially heightened transcriptional response to AD in these cell types in this M_e2x subgroup.

**Figure 1.**
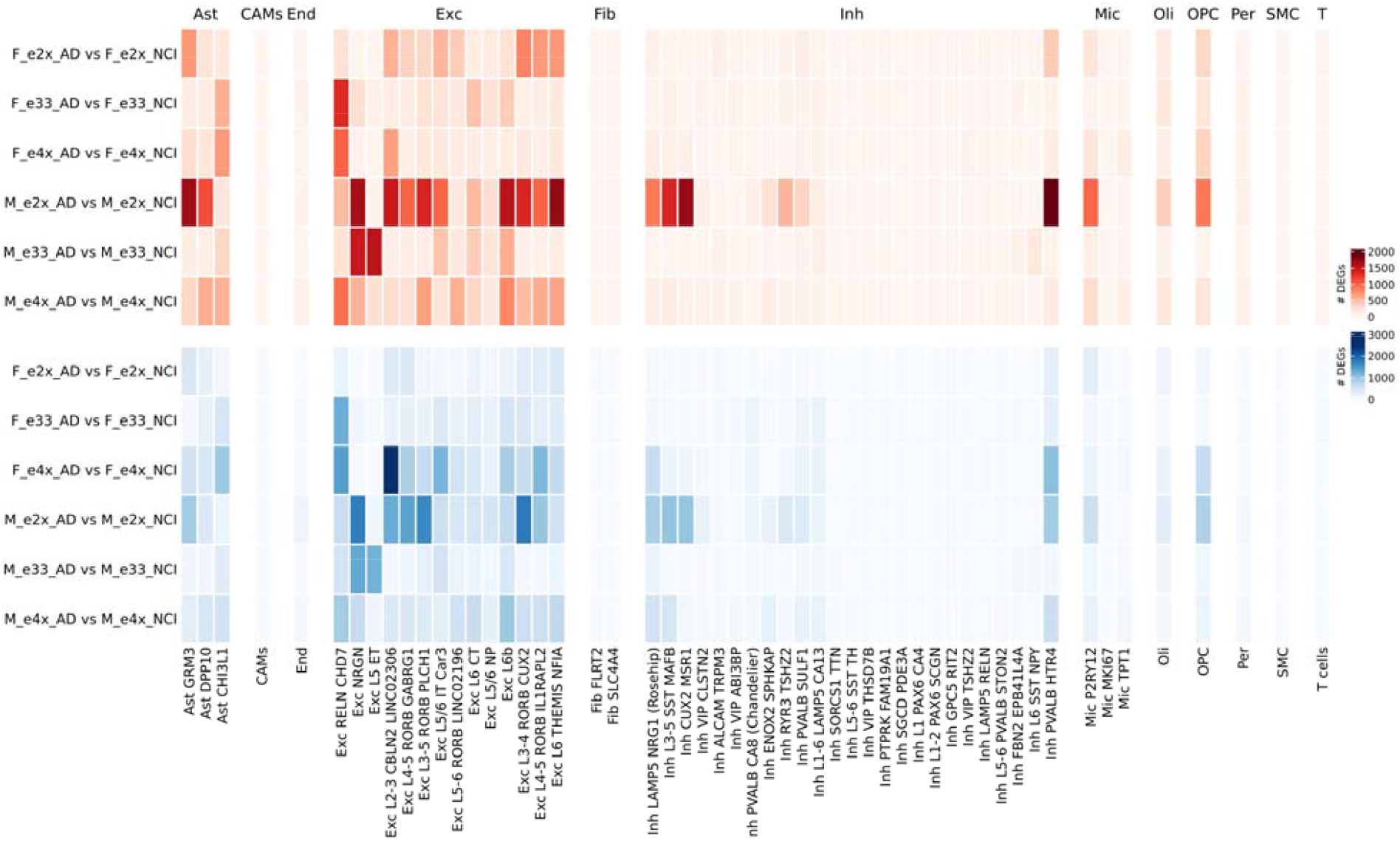
Heatmap showing the number of differentially expressed genes (DEGs) between AD samples and NCI samples within each sex-APOE group (F_e2x, F_e33, F_e4x, M_e2x, M_e33, M_e4x) across all the 54 cell clusters. The upper heatmap represents the number of upregulated DEGs (red), while the lower heatmap depicts the number of downregulated DEGs (blue). Each row corresponds to a specific comparison, and each column represents a distinct cell cluster.

### Comparative study of transcriptional signatures between different sex-APOE subgroups

To further explore transcriptional changes in AD, we conducted a series of pairwise comparisons to identify the overlap and divergence in differentially expressed genes (DEGs) across APOE genotypes and between sexes. These analyses aimed to identify both shared and unique transcriptional responses among subgroups, offering insights into how genotype and sex interact to influence molecular changes in AD in each brain cell type.

### Comparison of transcriptional signatures across *APOE* genotypes in females and males

We first compared the DEG signatures between females across *APOE* genotypes (F_e2x, F_e33, and F_e4x) to explore *APOE* genotype-specific transcriptional patterns in female (**Fig. 2A**). Comparisons included 1). F_e2x_AD vs. F_e2x_NCI and F_e33_AD vs. F_e33_NCI, 2). F_e2x_AD vs. F_e2x_NCI and F_e4x_AD vs. F_e4x_NCI, and 3). F_e4x_AD vs. F_e4x_NCI and F_e33_AD vs. F_e33_NCI. Notably, *APOE* e4 carriers showed more distinct transcriptional changes than the other genotypes, highlighting the unique impact of the e4 genotype on AD females.

**Figure 2.**
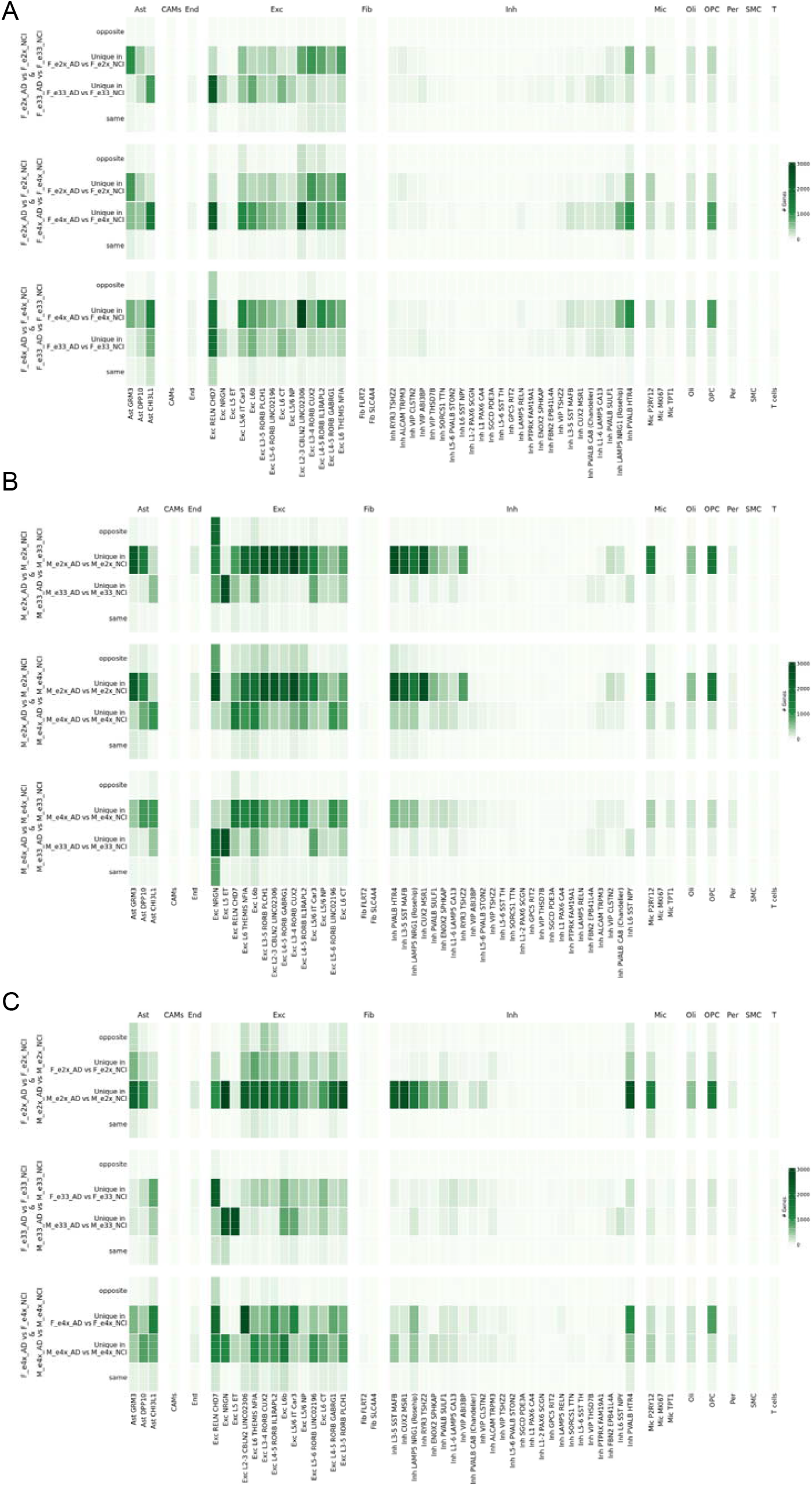
Comparative study of transcriptional signatures between different sex-APOE subgroups. **A)** Heatmap illustrating the overlap, unique, and opposite differentially expressed genes (DEGs) between female APOE groups (F_e2x, F_e33, and F_e4x) across cell clusters. Each row corresponds to a specific comparison, including: 1). F_e2x_AD vs. F_e2x_NCI and F_e33_AD vs. F_e33_NCI, 2). F_e2x_AD vs. F_e2x_NCI and F_e4x_AD vs. F_e4x_NCI, and 3). F_e4x_AD vs. F_e4x_NCI and F_e33_AD vs. F_e33_NCI. **B)** Heatmap illustrating the overlap, unique, and opposite differentially expressed genes (DEGs) between male APOE groups (M_e2x, M_e33, and M_e4x) across cell clusters. Each row corresponds to a specific comparison, including: 1). M_e2x_AD vs. M_e2x_NCI and M_e33_AD vs. M_e33_NCI, 2). M_e2x_AD vs. M_e2x_NCI and M_e4x_AD vs. M_e4x_NCI, and 3). M_e4x_AD vs. M_e4x_NCI and M_e33_AD vs. M_e33_NCI. **C)** Heatmap illustrating the overlap, unique, and opposite differentially expressed genes (DEGs) between males and females within each APOE genotype (e2x, e33, and e4x) across cell clusters. Each row corresponds to a specific comparison, including: 1). F_e2x_AD vs. F_e2x_NCI and M_e2x_AD vs. M_e2x_NCI, 2). F_e33_AD vs. F_e33_NCI and M_e33_AD vs. M_e33_NCI, and 3). F_e4x_AD vs. F_e4x_NCI and M_e4x_AD vs. M_e4x_NCI.

The same analysis was applied to the male *APOE* groups (M_e2x, M_e33, and M_e4x), including 1) M_e2x_AD vs. M_e2x_NCI and M_e33_AD vs. M_e33_NCI, 2) M_e2x_AD vs. M_e2x_NCI and M_e4x_AD vs. M_e4x_NCI, and 3) M_e4x_AD vs. M_e4x_NCI and M_e33_AD vs. M_e33_NCI. As shown in **Fig. 2B**, *APOE* e2 carriers exhibited a more pronounced transcriptional divergence from other genotypes, particularly within excitatory neurons, inhibitory neurons and astrocytes. This pattern suggests that *APOE* e2 carriers in males may activate certain distinct transcriptional programs in AD, with greater cell-type specificity than observed in females.

### Comparison Across Sexes Within Each *APOE* Genotype

Lastly, we examined the overlap and difference in DEGs between sexes for each *APOE* genotype (**Fig. 2C**). Comparisons included 1) F_e2x_AD vs. F_e2x_NCI and M_e2x_AD vs. M_e2x_NCI, 2) F_e33_AD vs. F_e33_NCI and M_e33_AD vs. M_e33_NCI, and 3) F_e4x_AD vs. F_e4x_NCI and M_e4x_AD vs. M_e4x_NCI. The results, visualized in **Figure 2C**, demonstrate significant sex-specific transcriptional differences in *APOE* e2 and *APOE* e4 genotypes. *APOE* e33 groups (F_e33_AD vs. F_e33_NCI and M_e33_AD vs. M_e33_NCI) had relatively fewer transcriptional differences between sexes across most cell clusters compared to e2x or e4x carriers. This observation aligns with the known biological role of the e33 genotype, which is generally considered neutral in terms of AD risk, in contrast to e2, which is protective, and e4, which is associated with heightened risk. The limited sex-specific divergence in e33 carriers suggests a more stable transcriptional landscape, potentially reflecting the absence of strong protective or risk-associated pressures on gene expression within this genotype.

### Similarity measures reveal shared and divergent transcriptional changes across sexes and *APOE* genotypes in AD

To quantitatively evaluate the similarity of differential expression patterns across sexes and *APOE* genotypes, we developed a new similarity measure for ternary vectors, termed as Zhang-Yu similarity measure (see **the Materials and Methods section** for the details). Each element in a ternary vector represents the differential expression status of a gene from a specific comparison with −1 standing for significant down-regulation, 0 for no significant change, and +1 for significant upregulation. The Zhang-Yu similarity measure quantitatively assesses the overlap, divergence, and directional consistency of gene expression changes across comparisons. Higher similarity scores indicate greater similarity while lower scores reflect divergence.

We employed this similarity measure to compare transcriptional patterns between females and males, incorporating results across all *APOE* genotypes (e2x, e33, and e4x) and all cell subclusters (**Figure 3A**). This allowed us to assess the consistency of sex-specific transcriptional responses across AD and NCI samples. Next, we extended the analysis to genotype-based comparisons, examining the similarity of transcriptional responses between *APOE* e2x and e33 carriers, as well as between e4x and e33 carriers (**Figure 3B, 3C**). For these analyses, we combined results across sexes and cell subclusters to identify genotype-specific patterns of differential expression.

**Figure 3.**
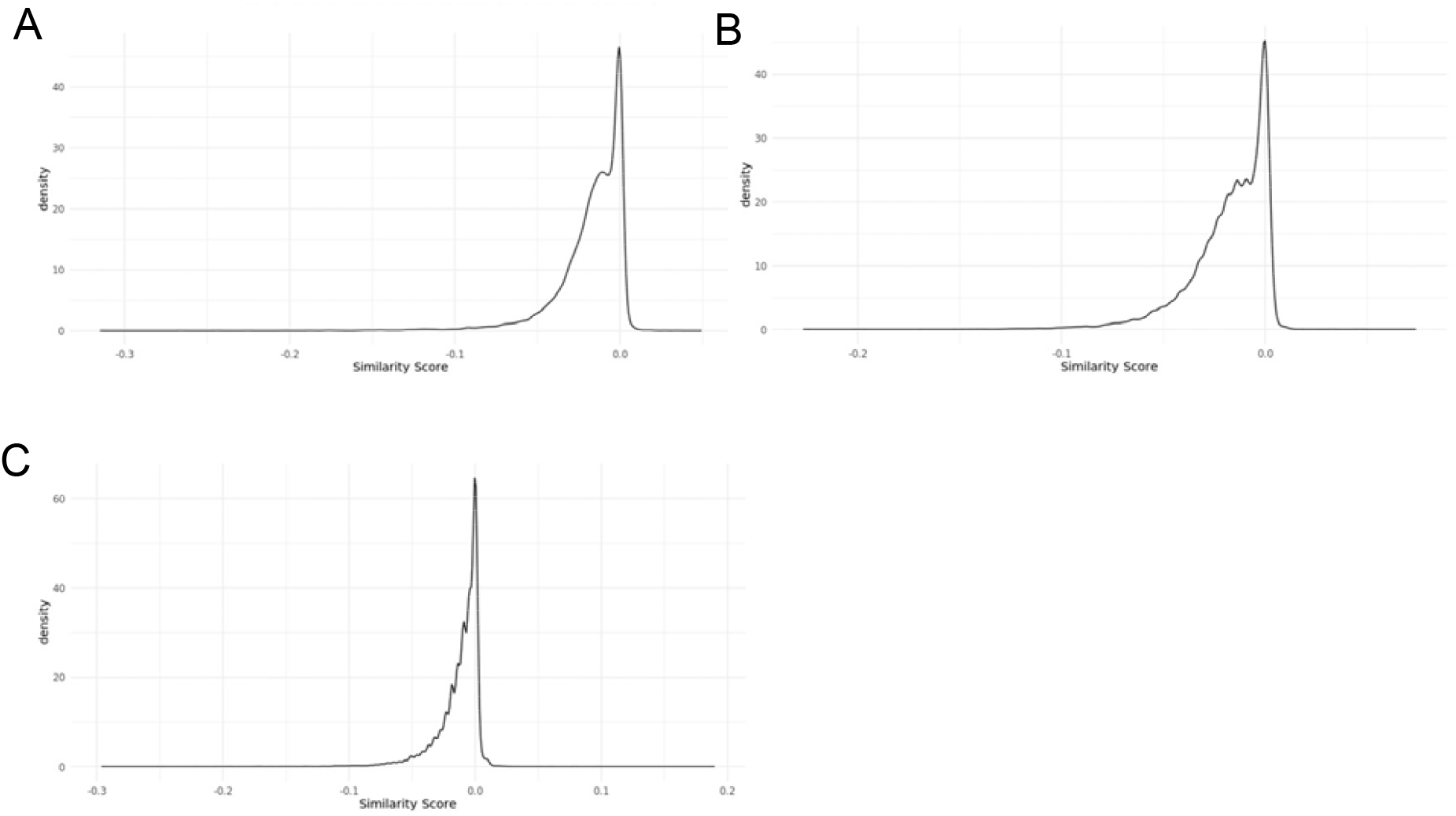
Distribution of similarity scores for different comparisons. **A)** The distribution of similarity scores for the comparison of females and males across all APOE genotypes (e2x, e33, and e4x). **B)** The distribution of similarity scores for the comparison of APOE e2x and e33 genotypes across both sexes. **C)** The distribution of similarity scores for the comparison of APOE e4x and e33 genotypes across both sexes.

To evaluate the effectiveness of the similarity metric, we plotted the expression patterns of approximately 500 genes, sampled evenly across the range of similarity scores, incorporating transcriptional changes associated with AD versus NCI across all *APOE* genotypes (e2x, e33, and e4x) and cell clusters for both females and males (**Figure 4**). For each gene, we analyzed the frequency of all possible combinations of expression patterns between females and males, such as consistent upregulation, consistent downregulation, or divergent patterns. As shown in the plot, genes with lower similarity scores are predominantly characterized by “opposite” and “different” cases, indicating divergent transcriptional responses between sexes. In contrast, genes with higher similarity scores tend to show more “same” direction cases, reflecting consistent transcriptional changes between females and males. This analysis demonstrates that the similarity metric effectively distinguishes shared versus divergent transcriptional patterns, supporting its utility in identifying sex- and genotype-specific transcriptional responses in Alzheimer’s disease.

**Figure 4.**
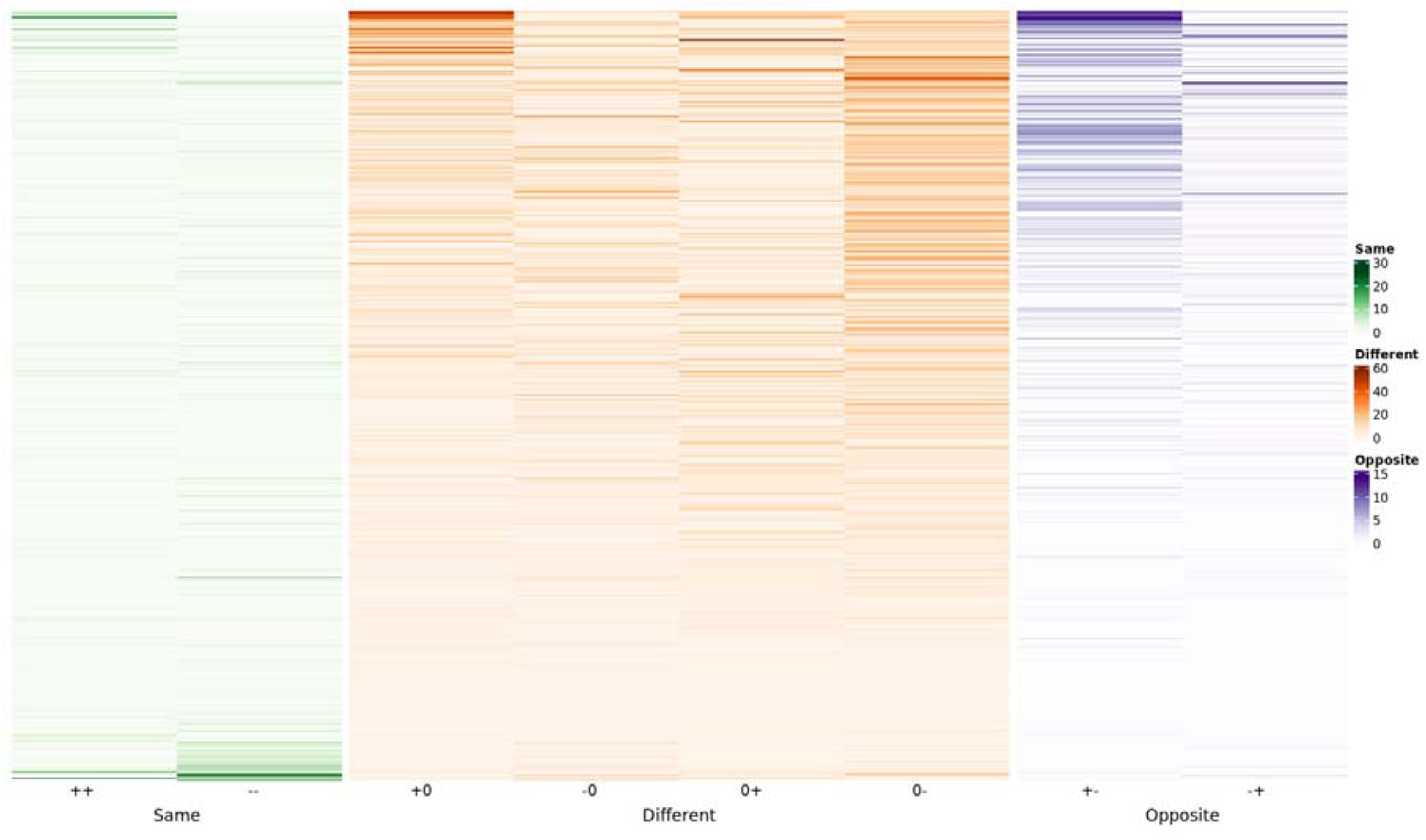
Heatmap illustrating the a set of approximately 500 genes that are evenly distributed across the full range of similarity scores for the comparison of differential expression patterns between females and males across all APOE genotypes. The heatmap categorizes gene expression changes as “Same” (++, --), “Different” (+0, 0+, -0, 0-), or “Opposite” (+-, -+), with color intensities indicating the number of occurrences across all cell subclusters.

Using the similarity score, we identified the top 25 (with the highest similarity) and bottom 25 (with the lowest similarity) genes for each of the three comparisons of our interest: sex differences, *APOE* e2x vs e33, and *APOE* e4x vs e33. For sex differences, the bottom 25 genes included *CLU* (Clusterin), a gene associated with AD risk and neuroinflammation, and *HSPB1* (Heat Shock Protein Beta-1) which is associated with stress responses in neurodegeneration. Both genes show divergent expression patterns between sexes (**Figure 5A**). Pathway enrichment analysis of the top 200 and bottom 200 genes further revealed distinct biological processes associated with shared and divergent transcriptional responses (**Figure 5B**). The top 200 genes, enriched in pathways such as “dopamine metabolism” and “histidine metabolism” suggest conserved roles in cellular metabolism, and hormone signaling across sexes. In contrast, the bottom 200 genes, representing divergent transcriptional responses, were enriched in immune-related and stress response pathways, including “cellular responses to external stimuli” “adaptive immune system” and “nonsense-mediated decay”. These enriched pathways point to sex-specific differences in immune and stress-response mechanisms in AD, highlighting potential molecular targets for understanding sex-related variability in disease progression.

**Figure 5.**
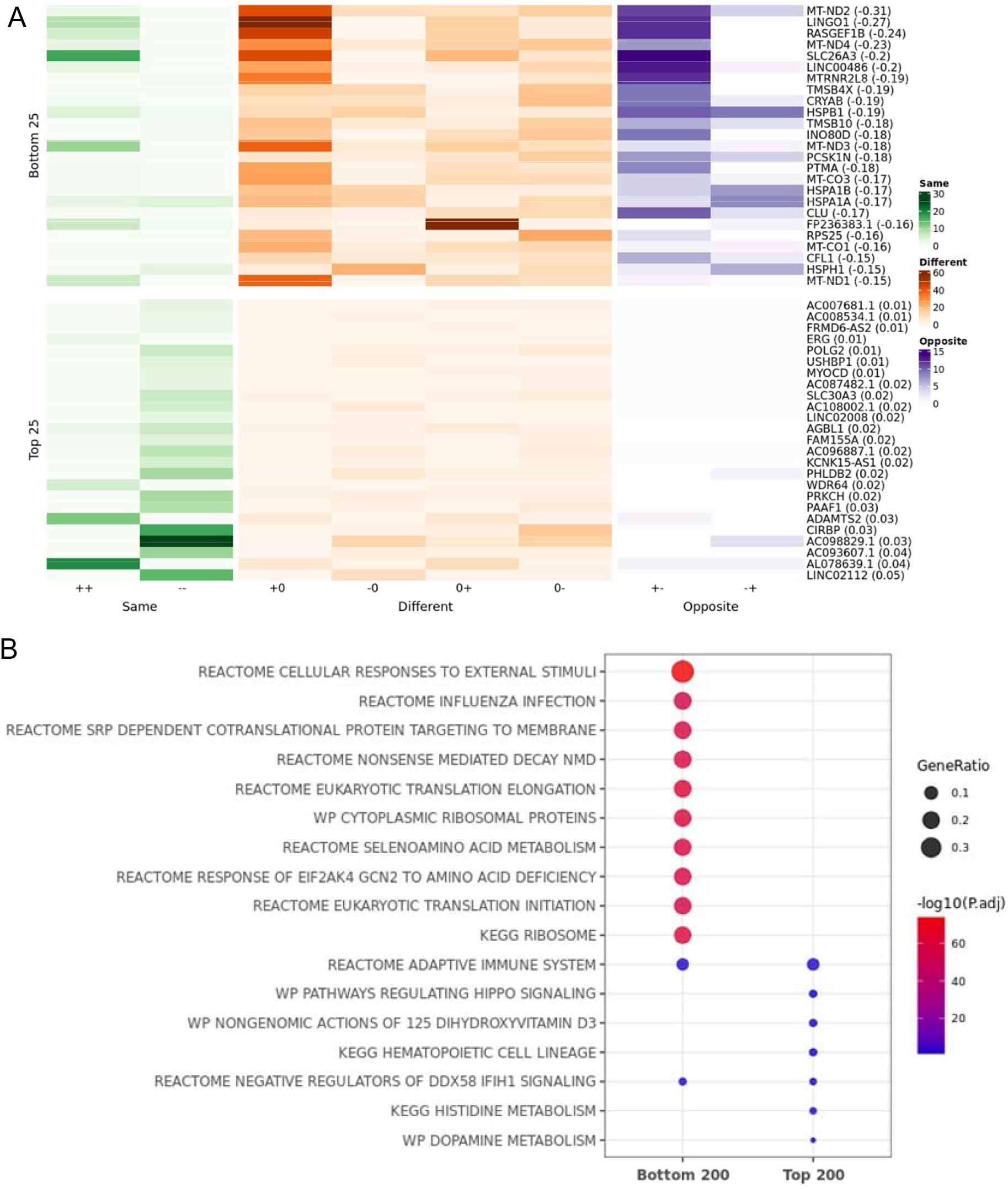
The top 25 genes displaying the greatest and least differences between sexes across APOE genotypes and cell types. **A)** Heatmap illustrating the top 25 and bottom 25 genes based on similarity scores for the comparison of differential expression patterns between females and males across all APOE genotypes and cell types. The heatmap categorizes gene expression changes as “Same” (++, --), “Different” (+0, 0+, -0, 0-), or “Opposite” (+-, -+), with color intensities indicating the number of occurrences across all cell subclusters. **B)** Pathway enrichment analysis for the top 200 and bottom 200 genes based on similarity scores in the comparison between females and males.

For the comparison between *APOE* e2x and *APOE* e33 genotypes, the top 25 genes with the highest similarity scores and the bottom 25 genes with the lowest similarity scores are shown in **Figure 6A**. Pathway enrichment analysis revealed that the top 200 genes were enriched in processes such as “fatty acid metabolism” and “SHC1 events in ERBB4 signaling,” emphasizing conserved roles in lipid metabolism and cellular signaling across these genotypes (**Figure 6B**). Conversely, the bottom 200 genes were enriched in pathways associated with translational regulation, such as “cytoplasmic ribosomal proteins” and “eukaryotic translation elongation,” as well as stress-response pathways like “cellular responses to external stimuli.” These enrichments highlight genotype-specific responses that may contribute to the differential impact of *APOE* e2 and *APOE* e33 in AD pathology. For the comparison between *APOE* e4x and *APOE* e33 genotypes, the top 25 genes with the highest similarity scores and the bottom 25 genes with the lowest similarity scores are shown in **Figure 7A**. Genes with the lowest similarity scores were enriched in mitochondrial function and energy metabolism pathways, including “oxidative phosphorylation” “electron transport chain” and “HSP90 chaperone cycle for steroid hormone receptors” (**Figure 7B**). These results suggest that mitochondrial processes and stress response pathways diverge significantly between *APOE* e4x and *APOE* e33 carriers.

**Figure 6.**
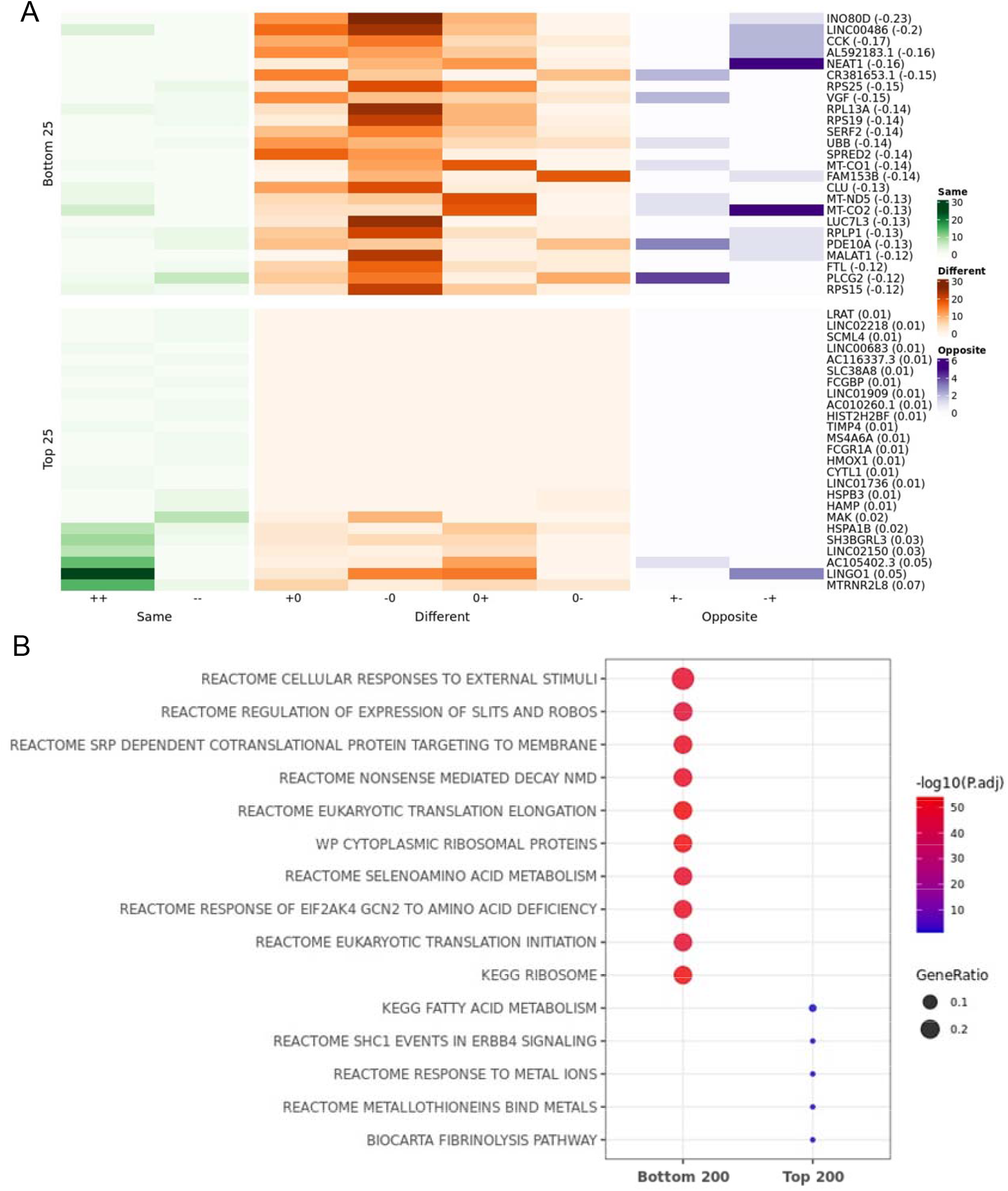
The top 25 genes displaying the greatest and least differences between APOE e2x and e33 carriers across sexes and cell types. **A)** Heatmap illustrating the top 25 and bottom 25 genes based on similarity scores for the comparison of differential expression patterns between APOE e2x and e33 carriers across sexes and cell types. The heatmap categorizes gene expression changes as “Same” (++, --), “Different” (+0, 0+, -0, 0-), or “Opposite” (+-, -+), with color intensities indicating the number of occurrences across all cell subclusters. **B)** Pathway enrichment analysis for the top 200 and bottom 200 genes based on similarity scores in the comparison between APOE e2x and e33 carriers.

**Figure 7.**
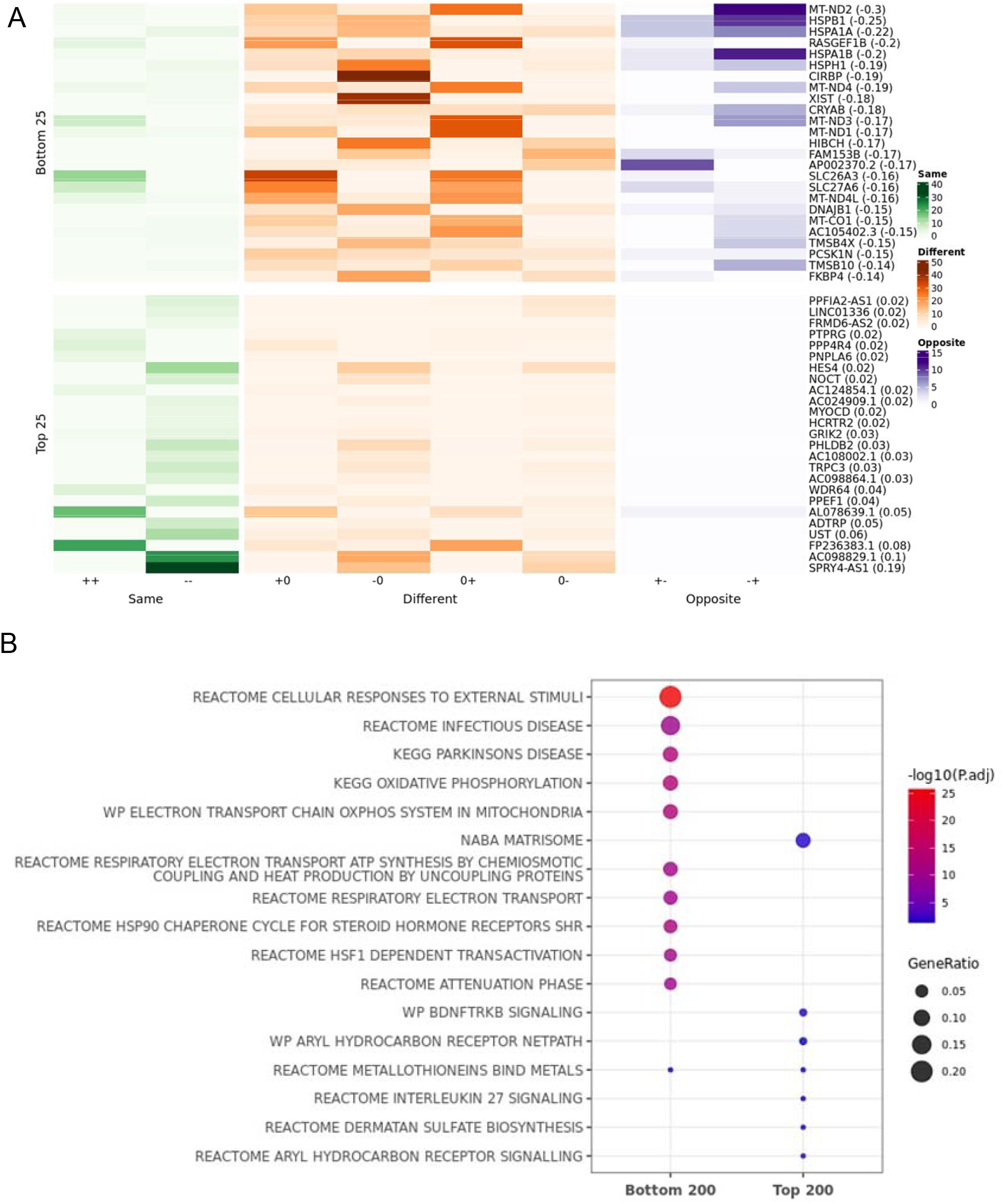
The top 25 genes displaying the greatest and least differences between APOE e4x and e33 carriers across sexes and cell types. **A)** Heatmap illustrating the top 25 and bottom 25 genes based on similarity scores for the comparison of differential expression patterns between APOE e4x and e33 carriers across sexes and cell types. The heatmap categorizes gene expression changes as “Same” (++, --), “Different” (+0, 0+, -0, 0-), or “Opposite” (+-, -+), with color intensities indicating the number of occurrences across all cell subclusters. **B)** Pathway enrichment analysis for the top 200 and bottom 200 genes based on similarity scores in the comparison between APOE e4x and e33 carriers.

## DISCUSSION

This study provides a comprehensive analysis of sex- and *APOE* genotype-specific transcriptional changes in AD using a snRNA-seq dataset from the ROSMAP cohort. By performing differential gene expression analysis across six stratified sex-*APOE* groups and applying a novel similarity metric, Zhang-Yule similarity measure, for ternary vectors, we uncovered shared and divergent transcriptional patterns across sexes and *APOE* genotypes. These findings highlight the interplay between genetic risk, sex, and cell-type-specific transcriptional responses in AD pathophysiology.

Male *APOE* e2 carriers exhibited the highest number of DEGs when comparing AD to NCI samples, suggesting a heightened transcriptional response in this subgroup. A majority of AD signatures were identified in excitatory neurons, inhibitory neurons, astrocytes, oligodendrocyte precursor cells, and one specific microglia cluster, highlighting the importance of these cell types in transcriptional responses to AD across all sex-*APOE* groups. Conversely, several cell clusters, including endothelial cells, CNS-associated macrophages (CAMs), pericytes, smooth muscle cells (SMCs), and T cells, showed no significant transcriptional changes, indicating a limited or negligible response to AD-related pathology in these populations.

In the comparison of transcriptional signatures across *APOE* genotypes in females, F_e4x AD signatures were notably divergent from those of F_e2x and F_e33. This divergence was particularly evident in comparisons such as F_e2x_AD vs. F_e2x_NCI and F_e4x_AD vs. F_e4x_NCI, and F_e4x_AD vs. F_e4x_NCI and F_e33_AD vs. F_e33_NCI, which displayed a higher number of DEGs that were unique to one condition. These unique DEGs suggest distinct transcriptional responses in F_e4x carriers, aligning with the heightened AD risk associated with *APOE* e4. Similarly, in males, M_e2x AD signatures were significantly divergent from M_e4x and M_e33 AD signatures. These findings highlight the distinct transcriptional responses associated with *APOE* e4 in females and *APOE* e2 in males, reflecting genotype-specific roles in AD pathology.

The use of the Zhang-Yule similarity measure allowed us to quantify shared and divergent transcriptional patterns across groups. Genes with high similarity scores represented conserved transcriptional responses, reflecting core molecular processes shared across sexes and genotypes. Conversely, genes with low similarity scores highlighted subgroup-specific pathways, particularly in immune responses and stress-related mechanisms. These patterns reinforce the complexity of AD pathology and emphasize the importance of examining both shared and divergent molecular changes in understanding the disease.

Overall, this study highlights the intricate interplay of sex and *APOE* genotype in shaping transcriptional responses in AD. By integrating cell-cluster-specific analyses with a similarity scoring approach, we provide a detailed view of the molecular heterogeneity underlying AD. These findings pave the way for future research to further elucidate the functional implications of identified pathways and develop precision medicine strategies that address the diverse molecular profiles of AD patients.

## MATERIALS AND METHODS

To investigate the genes, pathways, and cell types underlying sex and *APOE* isoform differences in AD pathology, we leveraged one of the most comprehensive single-nucleus RNA sequencing (snRNA-seq) datasets in AD research by Mathys et al1. This dataset includes 2.3 million nuclei isolated from the prefrontal cortex of 427 participants in the ROSMAP cohort, spanning a wide range of AD progression.

We used preprocessed read count data from the original study, which identified 54 high-resolution cell types grouped into 12 major categories. These include 14 excitatory neuron subtypes (Exc), 25 inhibitory neuron subtypes (Inh), oligodendrocytes (Oli), oligodendrocyte precursor cells (OPCs), 3 astrocyte subtypes (Ast), and 5 immune cell types (microglia (Mic), CNS-associated macrophages (CAMs), and T cells). The dataset also encompasses several vascular cell types, including endothelial cells (End), smooth muscle cells (SMCs), fibroblasts (Fib), and pericytes (Per).

### Cell cluster specific differential expression analysis

Differential expression analysis was performed for each sex-*APOE* group, including F_e2x, F_e33, F_e4x, M_e2x, M_e33, and M_e4x, comparing individuals with Alzheimer’s disease (AD) to control samples with no cognitive impairment (NCI). This analysis was conducted at the cell-cluster level to account for cell-type-specific transcriptional changes. We utilized the FindMarkers function in Seurat version 5 for cluster-specific DEG identification. Genes were included in the analysis if they were expressed in at least 10% of the cells in either group (min.pct = 0.1). To adjust for potential confounding factors, we incorporated covariates including nCount_RNA (total RNA counts), post-mortem interval (PMI), and age at death. The Benjamini-Hochberg (BH) method was used to adjust p-values for multiple testing. Significant differentially expressed genes (DEGs) were defined as those meeting the following thresholds: adjusted p-value (adj.p) < 0.05 and absolute fold change (|FC|) > 1.3. Filtered genes that passed these criteria were included in downstream analyses. Enrichment analysis was performed by using R package GOtest (V1.0.9).

### Similarity measure for evaluating similarity of transcriptional patterns between sex and *APOE* genotype subgroups

To quantitatively evaluate the similarity of differential expression patterns across sexes and *APOE* genotypes, we developed and applied a novel similarity measure termed Zhang-Yu similarity measure. This similarity measure calculates the similarity between two N-dimensional ternary vectors, i.e., ***X*** and ***Y***, representing differential expression statuses (AD versus NCI) of the genes in two different groups. In a ternary vector where each element takes three discrete values including -1, 0 and 1, each dimension represents the differential expression status of a gene as either up-regulation (+1), down-regulation (−1), or no significant change (0). The similarity score for two ternary vectors is defined as:

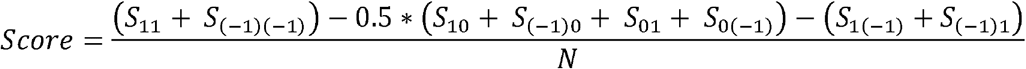

where, *S*_*ij*_ represents the number of occurrences of ***X(k)*** = *i* and ***Y(k) =*** *j*, where *k=1, …, N*. Therefore *S*_*11*_ represents for the number of the up-regulation matches in both groups, *S*_*(−1)(−1)*_ for the number of the down-regulation matches in both groups, and *S*_*10*_ for the number of mismatches with the upregulation in ***X*** and no change in ***Y***, and so on so forth. N is the number of genes evaluated in the differential expression analysis.

We applied this similarity measure to three key comparisons:

1. **Female vs. Male (F vs. M):**
  ◯ **X:** Differential expression results ({−1,0,1}) for females across all cell clusters and *APOE* genotypes: F_e2x_AD vs. F_e2x_NCI, F_e33_AD vs. F_e33_NCI, and F_e4x_AD vs. F_e4x_NCI.
  ◯ **Y:** Differential expression results ({−1,0,1}) for males across all cell clusters and *APOE* genotypes: M_e2x_AD vs. M_e2x_NCI, M_e33_AD vs. M_e33_NCI, and M_e4x_AD vs. M_e4x_NCI.
  ◯ **N:** Total number of cell clusters (54) multiplied by 3 comparisons = 162.
2. ***APOE* e2x vs. e33:**
  ◯ **X:** Differential expression results ({−1,0,1}) across all cell clusters for *APOE* e2x: F_e2x_AD vs. F_e2x_NCI and M_e2x_AD vs. M_e2x_NCI.
  ◯ **Y:** Differential expression results ({−1,0,1}) across all cell clusters for *APOE* e33: F_e33_AD vs. F_e33_NCI and M_e33_AD vs. M_e33_NCI.
  ◯ **N:** Total number of cell clusters (54) multiplied by 2 comparisons = 108.
3. ***APOE* e4x vs. e33:**
  ◯ **X:** Differential expression results ({−1,0,1}) across all cell clusters for *APOE* e4x: F_e4x_AD vs. F_e4x_NCI and M_e4x_AD vs. M_e4x_NCI.
  ◯ **Y:** Differential expression results ({−1,0,1}) across all cell clusters for *APOE* e33: F_e33_AD vs. F_e33_NCI and M_e33_AD vs. M_e33_NCI.
  ◯ **N:** Total number of cell clusters (54) multiplied by 2 comparisons = 108.

